# Decoupling T Cell Cytotoxicity: A CCL21+ICAM1-Based Synthetic Immune Niche Enhances Tumor Elimination by Accelerating Lytic Hit Delivery

**DOI:** 10.64898/2026.06.03.729495

**Authors:** Manish Kumar Gupta, Rawan Zoabi, Sofi Yado, Jörn Starruß, Benjamin Geiger, Haralampos Hatzikirou

## Abstract

Preserving T cell cytotoxic function during *ex vivo* expansion remains a major challenge for adoptive cancer immunotherapy. A synthetic immune niche (SIN) composed of immobilized CCL21 and ICAM1 was shown to improve T cell expansion while preserving cytotoxicity; however, it remains unclear which specific step of the T cell killing process is enhanced by the SIN stimulation. Here, we combined advanced imaging based on time-lapse microscopy with a mean-field model to analyze the distinct stages in killing of B16 melanoma cells by CD8^+^ T cells. This framework enabled us to resolve the tumor cell killing process into discrete steps throughout target cell engagement with SIN-treated T cells and lytic hit delivery. We found that tumor cell death is best explained by a multi-hit process, requiring approximately four discrete hits to trigger cell death. Model-based analysis identified an increase in the lytic hit delivery as the parameter that best accounts for the enhanced cytotoxicity of SIN-treated T cells, a difference not explained by changes in target encounter frequency or conjugate stability. Global sensitivity analysis further showed that tumor control is more strongly improved by enhancing lytic hit delivery than by increasing target encounter rates. These findings fundamentally reorient our understanding of optimized T cell manufacturing, suggesting that the lytic execution step may be the primary rate-limiting bottleneck in this in-vitro system, and that engineering strategies targeting granule polarization or discharge warrant prioritization alongside affinity-enhancement approaches.

## 1 INTRODUCTION

Adoptive cancer immunotherapy by tumor-infiltrating lymphocyte (TIL) and chimeric antigen receptor (CAR) T has demonstrated remarkable efficacy in refractory malignancies^1,2^, yet realizing its full potential remains a major challenge. A central manufacturing bottleneck is the *ex vivo* expansion of patient-derived T cells to clinically relevant numbers while preserving functional potency.^3^ Standard protocols rely on artificial APC surrogates such as anti-CD3/CD28-coated microbeads, which provide strong mitogenic stimulation but lack the structural and soluble complexity of the native lymphoid niche. As a result, T cells expanded under these conditions frequently differentiate toward terminal effector states and acquire features of exhaustion that compromise *in vivo* persistence and cytotoxic function.^4–7^ This proliferation–differentiation trade-off remains a pervasive obstacle in the field.

To address this limitation, a synthetic immune niche (SIN) was developed, which recreates, *ex vivo*, key stimulatory features of the lymphoid and the tumor-associated microenvironments.^8^ Substrates co-functionalized with the chemokine CCL21 and the integrin ligand ICAM1 support a robust T cell expansion while preserving, and in some cases augmenting, effector function.^9^ While the phenomenological outcomes of SIN treatment, namely, high yield of T cells with high killing capacity, are promising, the biological processes driving this enhanced cytotoxicity remain poorly understood. Enhanced killing can, in principle, be attributed to three distinct intercellular processes: An increased frequency of T cell–tumor cell encounters, prolonged contact duration, or a more efficient lytic hit delivery, once contact is established. Which of these processes SIN conditioning actually modulates — and to what degree — remains an open question with direct implications for therapeutic design. Distinguishing between these possibilities is challenging using standard population-level assays (e.g., Chromium-51 release or flow cytometry), which integrate these processes into a single endpoint measurement.

Mathematical modeling has long been used to study T cell cytotoxicity, but existing frameworks differ substantially in the degree to which they resolve the underlying biology. Early models of tumor–cytotoxic T lymphocyte (CTL) interactions typically used predator–prey or mass-action formulations, in which tumor cells proliferate and are eliminated through a single effective killing term proportional to effector and target abundance.^10,11^ While such models capture bulk killing kinetics, they do not distinguish between the elementary steps that generate cytotoxicity, such as target encounter, contact persistence, and productive lytic hit delivery. In contrast, stochastic and agent-based models, monitored by live-cell imaging, have shown that cytotoxicity is a multistep and heterogeneous process involving cell migration, conjugate formation, synapse maintenance, and serial or cooperative killing.^12–14^ These studies, together with single-cell experiments, further suggest that tumor cell death is often a complex, rather than binary process, and that may require repeated or additive sublethal hits rather than a single lethal event.^15–17^ However, although these richer models provide a detailed mechanistic view of the killing process, calibrating them from population-level measurements is often challenging because their larger parameter spaces can limit identifiability.

We therefore developed a coarse-grained mean-field model to address this biological question while retaining the key multistep structure. In this framework, target encounter is represented by an effective binding rate (*k*_1_), conjugate stability by a dissociation rate (*k*_2_), and productive killing by a lytic hit delivery rate (*k*_3_). By fitting the model to high-throughput live-cell imaging data of OT-I CD8^+^ T cells interacting with OVA/GFP-expressing B16 melanoma cells over 48 h (Fig 2), we investigated which of these effective kinetic steps differs between control and SIN-expanded T cells. We found that SIN-stimulated T cells show a marked increase in lytic hit delivery rate, a result we term “synaptic lethality.” Global sensitivity analysis further confirms that *k*_3_ exerts dominant control over tumor population dynamics, suggesting that future therapeutic optimization should prioritize the quality of the lytic event over the frequency or duration of cellular contacts.

**Figure 1.**
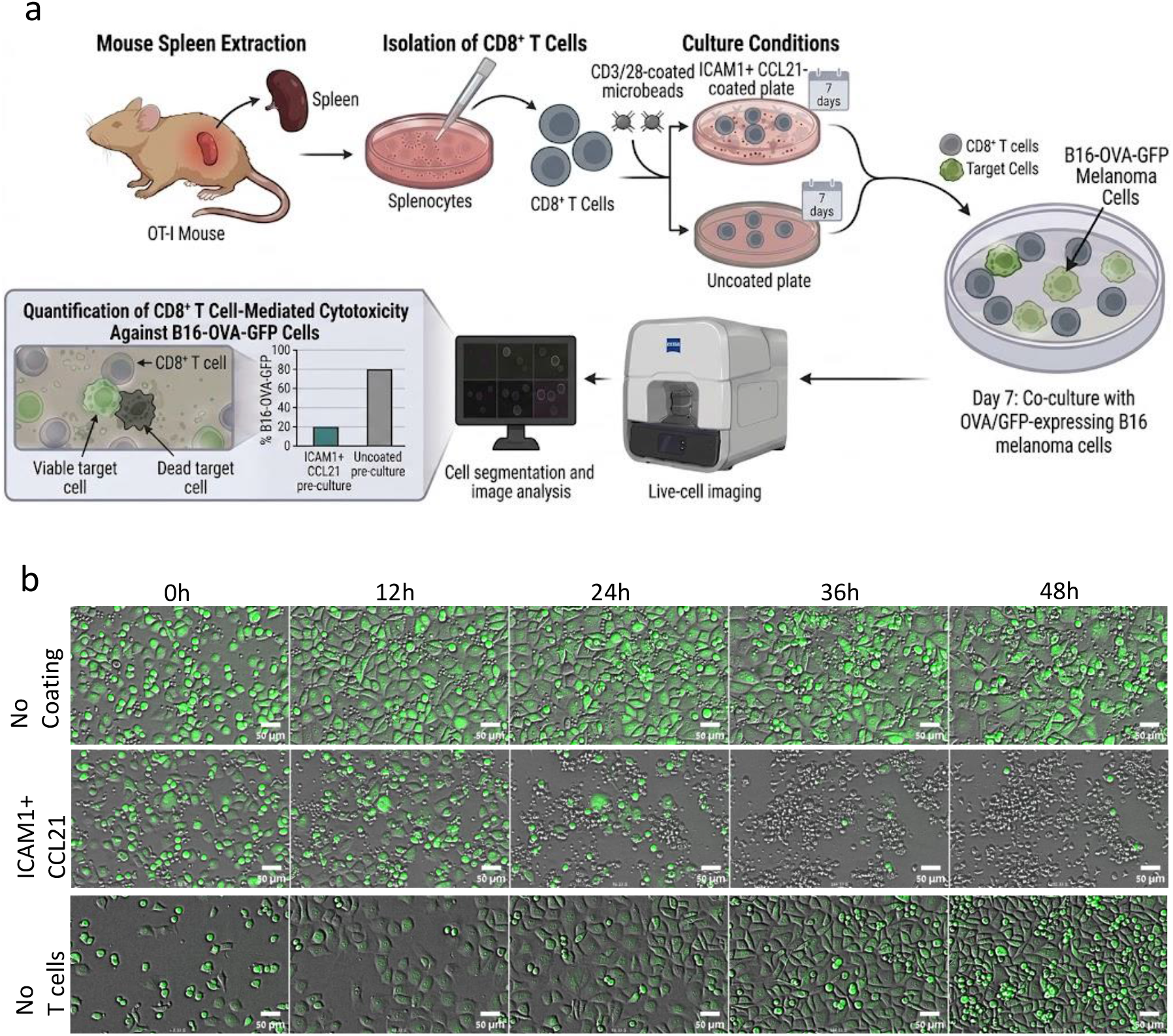
Schematic of the experimental workflow. (a) CD8^+^ T cells were isolated from spleens of OT-I mice, activated with CD3*/*28-coated microbeads, and cultured for seven days on either ICAM1 + CCL21-coated or uncoated surfaces. On day 7, SIN-treated and untreated T cells were co-cultured with antigen (OVA)- and GFP-expressing B16 melanoma cells. Live-cell imaging was performed using a Celldiscoverer 7 microscope, followed by cell segmentation and image analysis to quantify T-cell cytotoxicity, measured as loss of GFP fluorescence in target cells. (b) Representative transmitted-light and fluorescence time-lapse images of target cells co-cultured with control (untreated) T cells (top), SIN-treated T cells (middle), or without effector cells (bottom). Live target cells are identified by cytoplasmic GFP intensity (green). Live target cells are GFP^+^ (green); T cells are unlabeled. Scale bar, 50 *µ*m. Images are representative of n = 4 independent experiments, each performed in technical duplicate.

**Figure 2.**
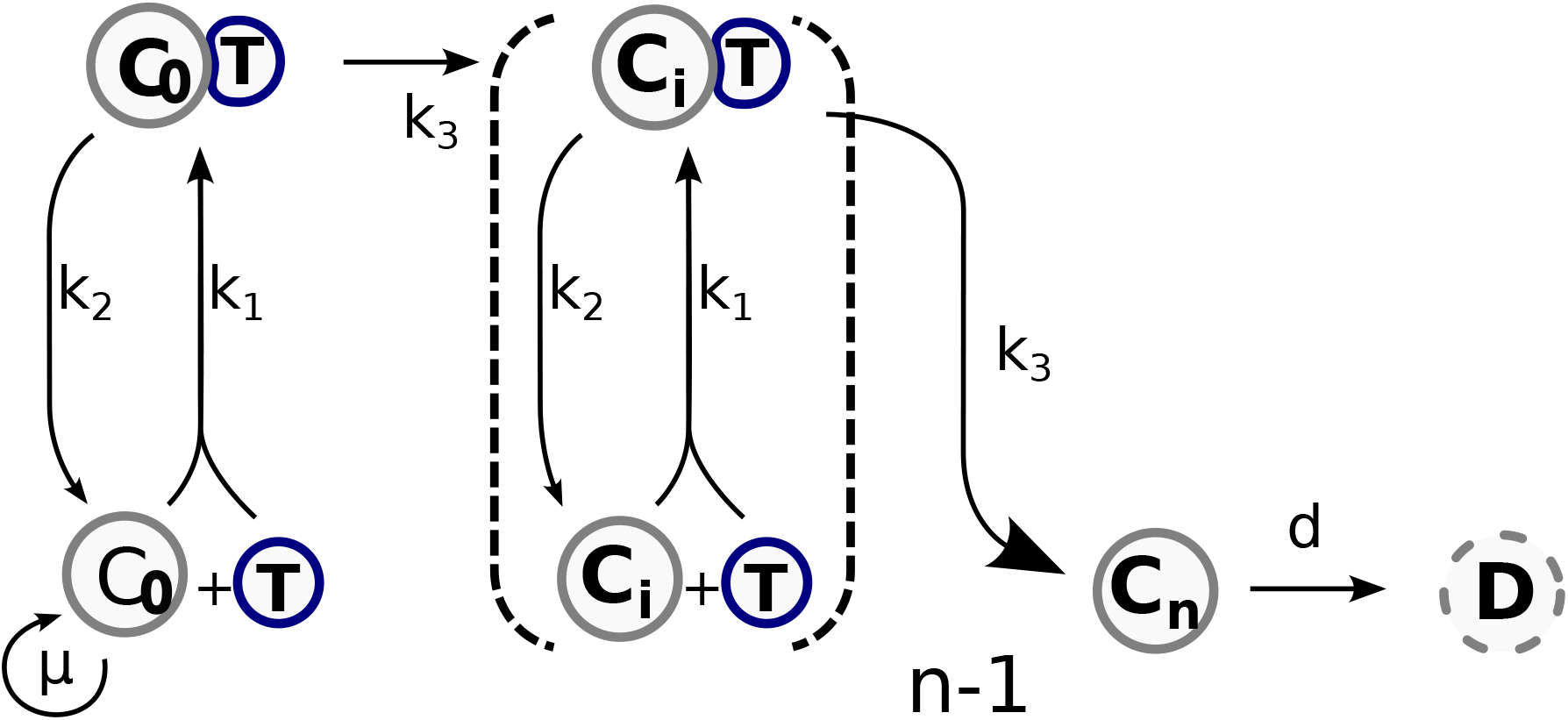
Schematic of the multi-hit cytotoxicity model. Tumor cells progress from the undamaged state *C*_0_ through successive damage states *C*_*i*_ upon T-cell-mediated lytic hit delivery. Free T cells bind targets with rate *k*_1_, dissociate with rate *k*_2_, and deliver hits within conjugates at rate *k*_3_. After *n* accumulated hits, target cells enter the apoptotic state *C*_*n*_ and decay with rate *d*.

**Figure 3.**
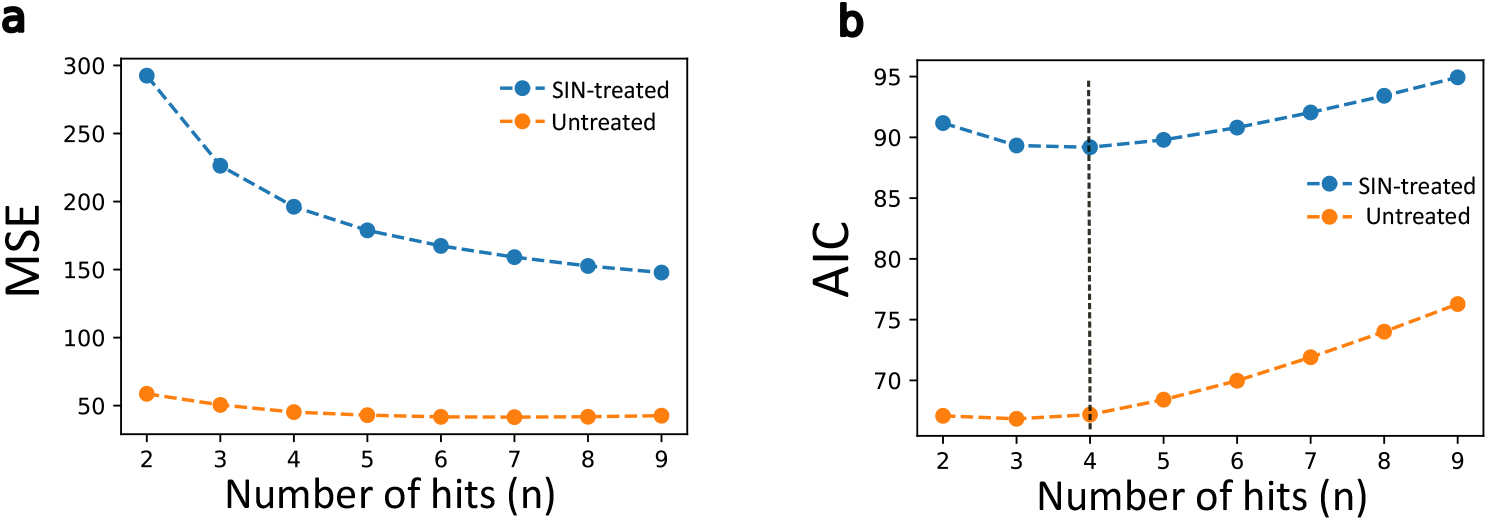
Model selection: (a) Mean residual square (MSE) obtained from model fitting plotted as a function of number of hits (*n*-hits) for both SIN-treated and untreated conditions. (b) Akaike Information Criterion (AIC) plotted as a function of number of hits (*n*-hits). The vertical dashed line indicates the number of hits selected from the model-selection analysis.

## 2 RESULTS

### A multi-hit framework accurately describes tumor elimination dynamics

To rigorously investigate the cytotoxic interplay between T cells and tumor cells, we constructed a mean-field mathematical model that moves beyond simple mass-action kinetics to incorporate the concept of cumulative damage. Experimental observations of the co-culture system, OT-I T cells and B16-OVA-GFP melanoma cells, revealed a nonlinear decay in the tumor cell population. Specifically, we observed an initial “shoulder” phase, representing a delay in the onset of significant tumor cell death, which is characteristic of systems where a threshold of damage must be exceeded before an irreversible state transition occurs.

We formulated the interaction as a system of coupled ordinary differential equations (ODEs) describing the transition of tumor cells through a series of discrete damage states, denoted by *C*_*i*_, where *i* represents the number of accumulated lethal “hits” (*i* = 0, 1, *…, n*). The model incorporates four fundamental biological processes:

1. Tumor growth: Viable undamaged tumor cells proliferate logistically with a specific growth rate *µ* and a carrying capacity *K*. We assume that partially damaged cells (*i >* 0) are arrested in the cell cycle, thus not contributing to proliferation.
2. Conjugation (Binding): Free T cells (*T*) bind to tumor cells in any state *C*_*i*_ with a second-order association rate constant *k*_1_, forming a physical complex representing the “search and capture” phase of cytotoxicity.
3. Complex dissociation (Unbinding): T cells dissociate from the complex with a first-order dissociation rate *k*_2_, returning the tumor cell to state *C*_*i*_ and the T cell to the free pool. This reversibility is crucial, as it allows T cells to detach and engage new targets (serial killing).
4. Hit delivery (Lethal step): Within the conjugated complex, the T cell executes a lytic event (e.g., granule exocytosis) with a first-order rate constant *k*_3_. This transition moves the tumor cell from damage state *C*_*i*_ to *C*_*i*+1_, effectively advancing it toward apoptosis.

The final state *C*_*n*_ represents the irreversibly committed apoptotic cell, which subsequently decays (loses membrane integrity and fluorescence) at a rate *d*. The mathematical representation of this system is given by the following set of equations:

The model further assumes that sublethal damage is not repaired over the 48 h experimental timescale, such that tumor cells progress irreversibly through the discrete damage states. This approximation is motivated by the high-affinity OT-I/B16-OVA system, in which cytotoxic damage is expected to accumulate on experimentally relevant timescales. For simplicity, we also neglect T-cell proliferation and exhaustion during the assay, and treat the free T-cell evolving only through binding and unbinding, while the total T-cell remains temporally constant.

### In-vitro B16 melanoma cells require approximately four hits to commit to apoptosis

A central question in T cell biology is the “quantification” of lethality, namely, is a single degranulation event sufficient to induce target cell death? To answer this, we treated the number of hits (*n*) as a key structural parameter. We fitted the model to time-series measurements of tumor population size under both SIN-treated and untreated conditions, systematically varying *n* from 1 (single-hit) to 9.

Model performance was evaluated using two metrics: the mean squared error (MSE), quantifying the deviation between the simulations and the data, and the Akaike information criterion (AIC)^18^, which penalizes model complexity and guards against overfitting. As expected, increasing *n* introduced a longer delay (shoulder) in the tumor decay curve, improving the fit to the early time points. The MSE decreased sharply as *n* increased from 1 to 3, and then plateaued. This indicates that single-hit kinetics (*n* = 1) are insufficient to capture the observed lag in B16 apoptosis.

The AIC provided a clearer criterion for model selection. For the SIN-treated condition, the AIC reached a distinct minimum at *n* = 4. For the untreated condition, the minimum was observed at *n* = 3, with *n* = 4 yielding a very similar value. To maintain structural consistency and enable direct parameter comparisons between conditions, we selected *n* = 4 as the consensus hit number for the remainder of the analysis. We note that this inferred hit number is specific to the model’s assumptions, in particular the irreversibility of sublethal damage and the uniformity of hit strength. This finding suggests that B16 melanoma cells function as signal integrators, requiring the accumulation of approximately four discrete cytotoxic hit events to reach an irreversible commitment to cell death.

### Direct experimental estimation constrains the parameter space

One of the central challenges in Systems Biology is parameter identifiability—the ability to uniquely determine parameter values from observed data. To mitigate this, we employed a “divide and conquer” strategy, estimating a subset of model parameters directly from image-derived measurements and then fitting only the remaining parameters using the ordinary differential equation (ODE) model, thereby reducing the number of free parameters.

### Decay rate (*d*)

We quantified the duration of the cell death phase as the time interval between cell rounding (morphological commitment) and the complete loss of GFP fluorescence (membrane permeabilization/death), with this combination serving as a proxy for target cell death. Tracking individual cell-death events yielded a distribution of decay time durations with a population mean *T*_avg_ ≈ 164 min (Fig. 4 b). Assuming a first-order (exponential) execution phase, the decay rate was estimated as 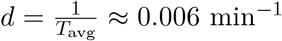. This value was consistent across experimental conditions, suggesting that the execution phase of death is intrinsic to the tumor cell and is not statistically significantly altered by the T cell’s activation history (SI figure 2).

**Figure 4.**
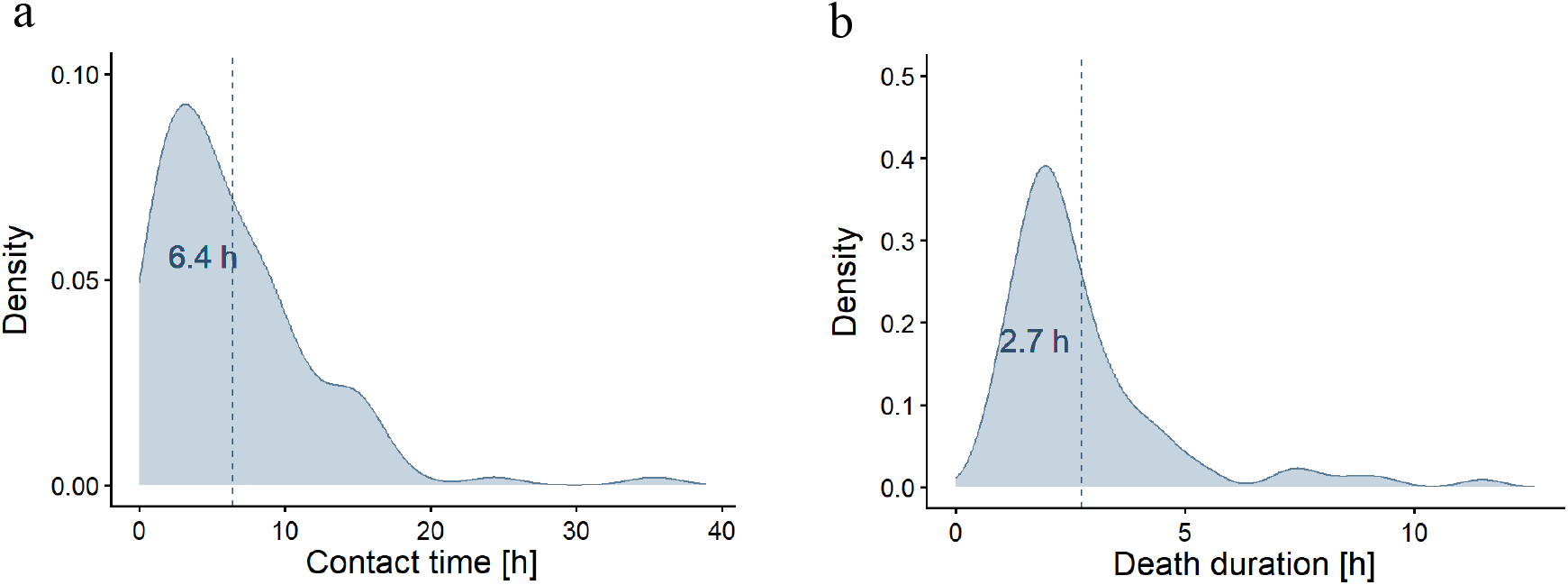
Experimental estimate of T cell-target cell dissociation rate (*k*_2_) and cell death duration. Density plots of the contact time between T cells and target cells (a) and the interval between target cell rounding up and the loss of GFP (cell death duration ; b), based on combined data from SIN-treated and untreated conditions. Vertical lines indicate the mean.

### Dissociation rate (*k*_2_)

Similarly, we measured the duration of physical conjugation events between T cells and tumor cells (Fig. 4 a). We assumed that T cell–tumor cell dissociation follows a first-order stochastic process with constant rate *k*_2_. Based on the measured mean attachment duration ⟨*T*_attach_⟩ ≈ 400 min ; the dissociation rate was estimated as 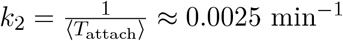. No measurable differences in attachment durations were detected between SIN-treated and untreated T cells (SI figure 2).

### Profile likelihood analysis reveals *k*_3_ as the driver of SIN-mediated cytotoxic enhancement

With *d* and *k*_2_ fixed by direct measurement, the core parameter estimation task focused on the three latent kinetic rates: the conjugation rate *k*_1_, the lytic hit delivery rate *k*_3_, and the proliferation rate *µ*. We used profile analysis to assess practical identifiability by fixing one parameter at a time while re-optimizing the remaining parameters by minimizing the mean squared errors (MSE) between model predictions and data. This mapped the MSE landscape and enabled the extraction of approximate 95% confidence intervals.

Initial attempts to fit the model to individual datasets resulted in broad, flat valleys in the likelihood profiles for *k*_1_ and *k*_3_, indicating practical non-identifiability^19^: The data were equally well explained by either a high-binding/low-killing regime or a low-binding/high-killing regime. To systematically investigate this limitation, we performed a simulation-based identifiability analysis using synthetic datasets generated from the model. Profile likelihood analysis^20^ of a single synthetic ROI confirmed that the parameters were not practically identifiable under an individual curve fitting strategy (SI Fig. 1). To resolve this ambiguity we adopted a co-fitting strategy, simultaneously fitting the model to multiple synthetic datasets derived from regions of interest (ROIs) with varying initial effector and target cell numbers while maintaining the same initial E: T ratio. This enriched dataset successfully constrained the parameter space, yielding well-defined U-shaped likelihood profiles for both parameters, indicative of practical identifiability (SI Fig. 1). Having established the data requirements for practical identifiability, we applied the co-fitting strategy to the experimental datasets for both SIN-treated and untreated conditions.

### Parameter comparison (SIN treated vs. untreated)

- **Conjugation rate (*k*_1_):** Analysis of the untreated condition revealed unexpected density-dependent effects. In high-density ROIs, untreated T cells appeared less effective at conjugating than predicted by mass action, likely arising from tumor cell clustering or spatial shielding. To account for this, we introduced a phenomenological prefactor *γ* applied to *k*_1_, such that the effective conjugation rate in high-density ROIs becomes *γk*_1_, correcting for the reduction in available tumor cell surface area due to clustering and spatial shielding. However, the introduction of this additional parameter came at a cost to identifiability: with *k*_1_ and *γ* jointly governing the effective conjugation rate, the available data cannot uniquely constrain *k*_1_ alone, as the same observed dynamics can be equally well explained by different combinations of *k*_1_ and *γ*. This resulted in a flat, unbounded likelihood profile for *k*_1_ in the untreated condition, in contrast to the treated condition where no such correction was necessary and *k*_1_ was successfully identified (Fig. 5 b and Fig. 6 c). Consequently, a statistically meaningful comparison of *k*_1_ between conditions cannot be made, and we conclude that the available data are insufficient to resolve potential differences in conjugation rate.
- **Hit Delivery rate (*k*_3_):** In contrast, the estimated hit delivery rate *k*_3_ showed a profound and statistically significant increase in SIN-treated cells, with value approximately 4 times the magnitude of their untreated counterparts (*p* = 0.0003; Table 1).
- **Proliferation rate (*µ*):** No statistically significant difference in *µ* was found between conditions (*p* = 0.06; Table 1), suggesting that SIN exposure does not substantially alter intrinsic tumor growth kinetics.

**Figure 5.**
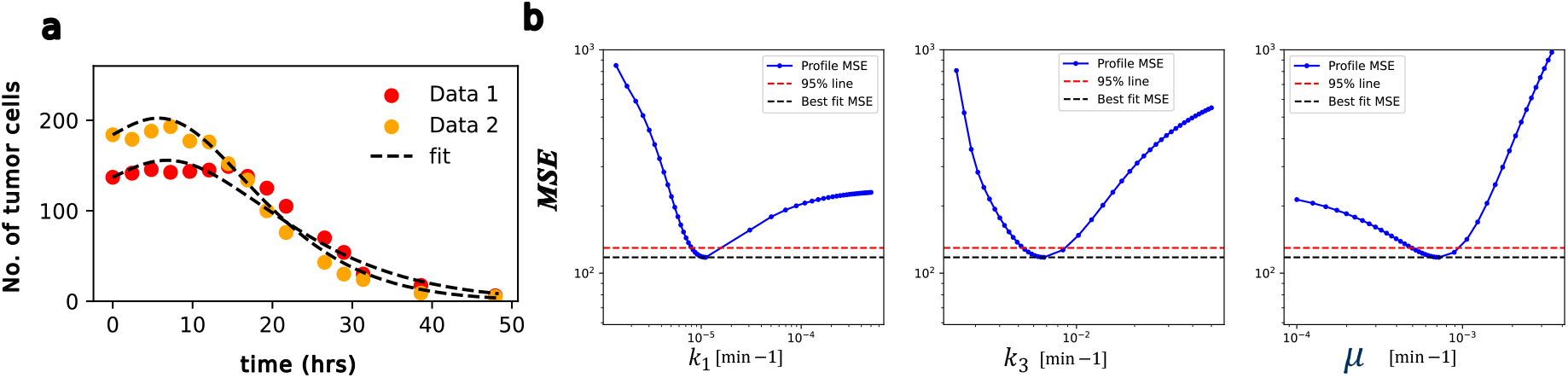
Parameter estimation for the SIN-treated condition: (a) Model fit to tumor-cell count trajectories measured from two independent regions of interest (Data 1 and Data 2) under the SIN-treated condition. The dashed black line shows the model prediction fitted jointly to the ROI-specific trajectories. (b) Profile-MSE curves for the fitted parameters *k*_1_, *k*_3_, and *µ*, used to assess practical parameter identifiability. The red dashed horizontal line indicates the 95% confidence threshold, and the confidence intervals are determined by the intersections of this threshold with the profile curves.

**Figure 6.**
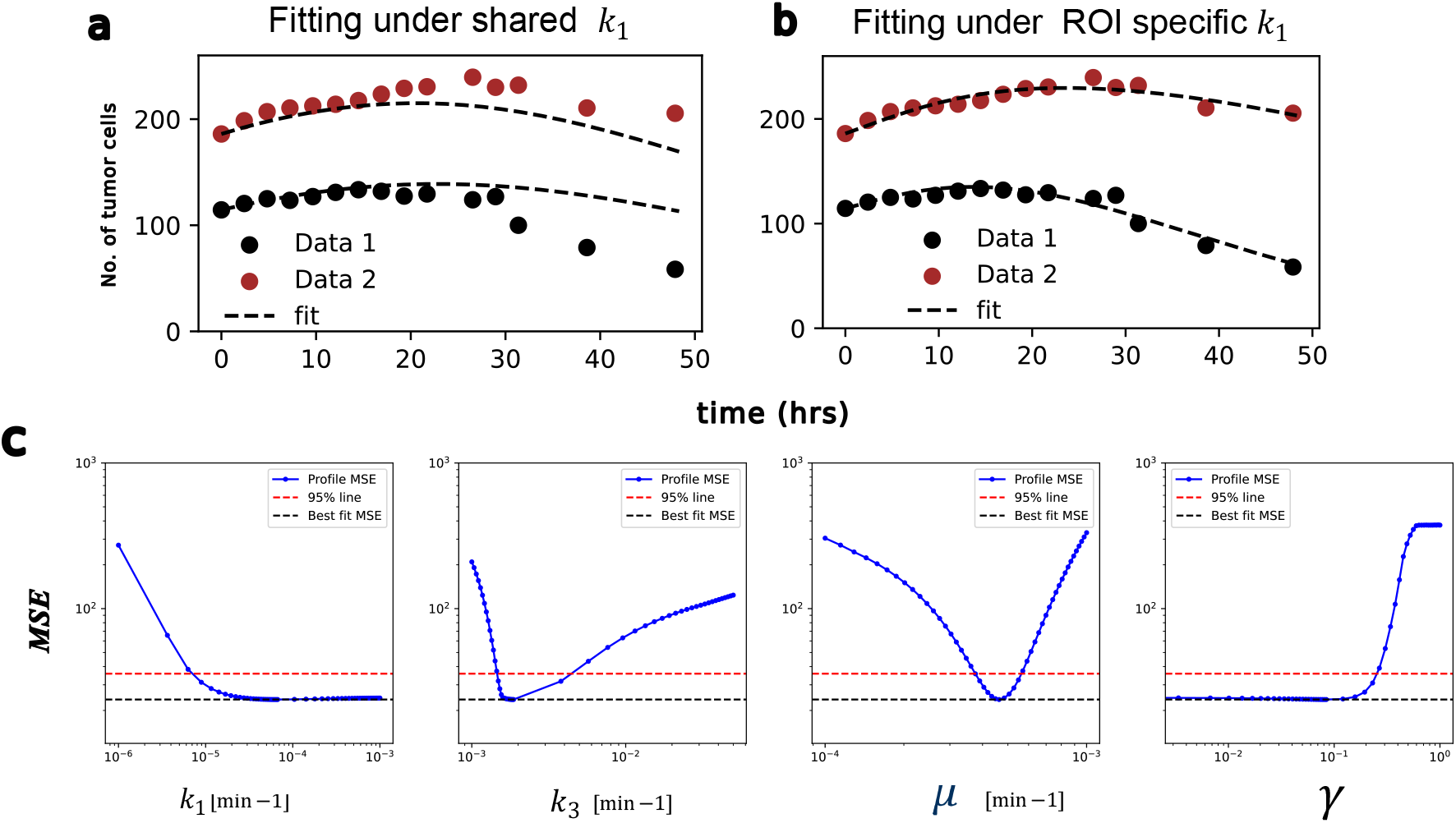
Parameter estimation for the untreated case: (a) Illustration of the model’s inability to adequately fit data collected from different ROIs using a shared set of parameter values, demonstrating poor agreement across datasets. (b) Improved fit achieved by allowing attachment rate *k*_1_ to vary between ROIs. (c) Practical identifiability test for the untreated case, showing *k*_1_ cannot be uniquely determined with the current dataset.

**Figure 7.**
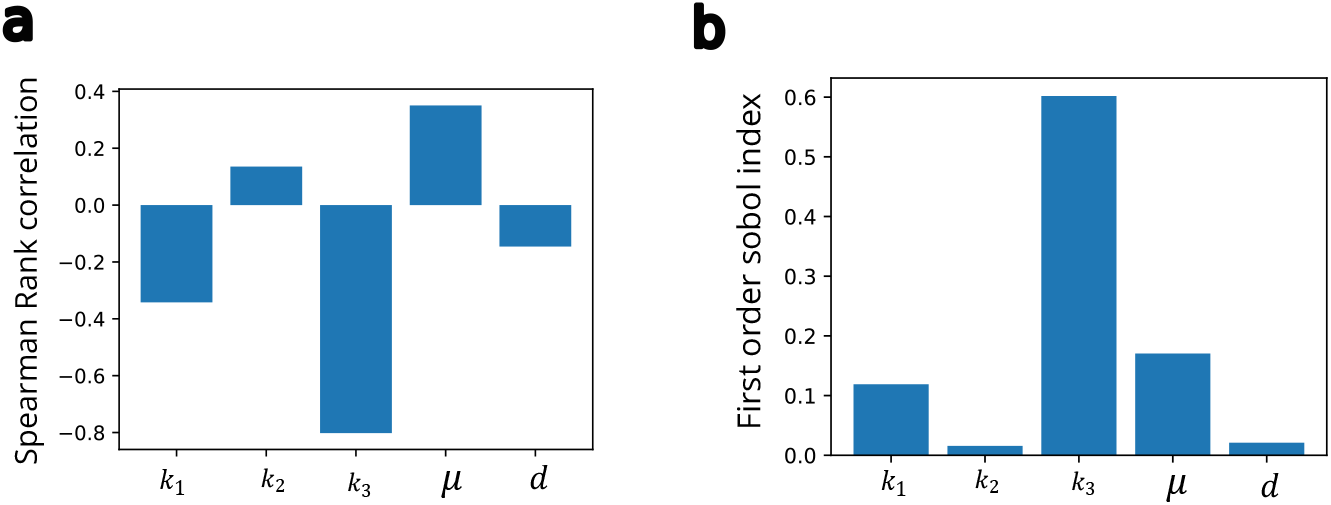
Global sensitivity analysis: (a) Spearman rank correlation coefficients for all parameters, indicating the direction of association with tumor burden. (b) First-order Sobol indices for each parameter, which quantify the contribution of each parameter to the overall variance of the model output, thereby providing insight into the relative importance of each parameter. Sensitivity analysis was performed by varying *k*_1_ over [5.5 ×10^*−*6^, 5.5 ×10^*−*5^], *k*_2_ and *k*_3_ over [10^*−*3^, 10^*−*2^], *µ* over [10^*−*4^, 10^*−*3^], and *d* over [10^*−*3^, 10^*−*2^].

**Table 1.** Estimated model parameters for SIN-treated and untreated conditions: Mean estimates are reported together with 95% confidence intervals (CI). All parameter values are given in min^*−*1^. The final column reports the *p*-value for the comparison between SIN-treated and untreated conditions. NA indicates that the upper confidence bound could not be determined from profile likelihood analysis, indicating practical non-identifiability of *k*_1_ under the current experimental conditions.

| Parameter | Treated | | Untreated | | $p$-value |
| --- | --- | --- | --- | --- | --- |
| | Mean ($\text{min}^{-1}$) | 95% CI | Mean ($\text{min}^{-1}$) | 95% CI | |
| $k_1$ | $1.1 \times 10^{-5}$ | $[0.8, 2] \times 10^{-5}$ | $6.7 \times 10^{-5}$ | NA | NA |
| $k_3$ | $6.7 \times 10^{-3}$ | $[5, 9] \times 10^{-3}$ | $1.8 \times 10^{-3}$ | $[1.5, 5] \times 10^{-3}$ | 0.0003 |
| $\mu$ | $7.2 \times 10^{-4}$ | $[5, 10] \times 10^{-4}$ | $4.62 \times 10^{-4}$ | $[3.7, 5.8] \times 10^{-4}$ | 0.06 |

**Table 2.** 95 % critical values for the difference Δ_𝒳_ 2

| $q$ profiled parameters | d.f. | $\chi_{q, 0.95}^2$ |
| --- | --- | --- |
| 1 | 1 | 3.84 |
| 2 | 2 | 5.99 |
| 3 | 3 | 7.81 |

This analysis yields the central finding of the study: the hit delivery rate *k*_3_ is approximately four-fold higher in SIN-treated T cells, while conjugate stability (*k*_2_) is statistically indistinguishable between conditions. We cannot make an equivalent comparative statement about encounter rate *k*_1_, because it remains practically non-identifiable for the untreated condition under the current experimental design. Its contribution could be resolved by encounter-rate measurements using single-cell tracking of T cell migration before conjugate formation. We therefore conclude specifically that accelerated lytic hit delivery is a sufficient and statistically supported mechanistic explanation for the enhanced cytotoxicity of SIN-treated T cells.

### Global sensitivity analysis identifies *k*_3_ as the primary determinant of cytotoxic efficacy

To validate the functional relevance of these findings, we performed a global sensitivity analysis to identify parameters that most strongly influence cytotoxic efficacy, defined as the cumulative tumor burden over 48 hours (time-integrated tumor population over 48 hours, AUC), where lower AUC values indicate improved tumor control.

We employed two complementary sensitivity metrics:

1. **Spearman rank correlation^21^:** This analysis assessed the direction and monotonicity of the relationship between each parameter and tumor burden. As expected, both *k*_1_ and *k*_3_ showed strong negative correlations with tumor AUC (i.e., increased binding or enhanced killing efficiency reduces tumor burden).
2. **Sobol indices^22^:** To quantify the magnitude of influence, we computed first-order Sobol indices, which decompose the variance of the model output into contributions from individual parameters.

The Sobol analysis revealed a clear hierarchy of influence. The hit delivery rate *k*_3_ emerged as the dominant parameter, explaining the largest fraction of the variance in tumor burden. The conjugation rate *k*_1_ was the second most influential factor, while the dissociation rate *k*_2_ and apoptosis rate *d* had negligible impacts on the outcome variance.

This dominance of *k*_3_ over *k*_1_ can be understood through the lens of the multi-hit bottleneck. In a regime where multiple hits are required (*n* = 4) for tumor cell elimination, a T cell that binds frequently (high *k*_1_) but delivers hits inefficiently (low *k*_3_) remains functionally suboptimal. Consequently, the most effective lever for improving tumor clearance is to maximize the lethality of each contact, rather than the frequency of contacts.

## 3 DISCUSSION

SIN engineered with the lymphoid cues CCL21 and ICAM-1 enhance the functional potency of adoptively expanded T cells, but the biophysical basis of this improvement has remained unresolved. In particular, it has been unclear whether the enhanced cytotoxicity of CCL21/ICAM-1-treated T cells reflects improved target encounter, greater conjugate stability, or more efficient lethal hit delivery after synapse formation. By integrating long-term live-cell imaging with a mechanistic mean-field model, we were able to separate these alternatives at the level of effective kinetic parameters. Our work shows that the dominant effect of CCL21/ICAM-1-based SIN is not on target engagement itself, but on lytic progression within the immune synapse.

Our model suggests the lytic hit delivery rate (*k*_3_) as the key parameter enhanced by SIN stimulation. The “hit” in our model represents the aggregate of the molecular events downstream of synapse formation, including Microtubule-Organizing Center (MTOC) polarization toward the synaptic interface, transport of lytic granules along microtubules, granule fusion with the plasma membrane, and diffusion of perforin and granzymes across the synaptic cleft^23^. Recent mechanistic studies provide plausible pathways by which substrate-bound ICAM-1 could specifically accelerate these steps. Previous work has shown that CCR7 and LFA-1 signaling can amplify synapse-associated signaling and cytoskeletal organization in T cells.^24–26^ Consistent with this, CCL21 + ICAM1 stimulation may prime T cells for accelerated lytic execution without necessarily altering the duration of the T cell–tumor conjugate–a phenotype we term *“synaptic lethality hypothesis”*.

The sensitivity analysis identifying lytic hit delivery rate (*k*_3_) as the primary parameter influencing cytotoxic efficacy provides conceptual guidance for T cell engineering strategies. Current approaches often focus on increasing CAR or TCR affinity to enhance tumor recognition (*k*_1_). However, our results suggest that such strategies may yield diminishing returns, particularly in the context of solid tumors requiring multiple hits. If *k*_3_ is limiting, increasing affinity may instead prolong non-productive T cell–target interactions. Therapeutic optimization should therefore focus on molecular determinants governing *k*_3_: This could involve:

- **Co-stimulatory Domain Optimization:** Selecting CAR intracellular domains (e.g., CD28 vs. 4-1BB) that specifically accelerate granule polarization dynamics.^27,28^
- **Granule Loading:** Genetic overexpression of Granzyme B or Perforin to ensure that each degranulation event is more potent (effectively increasing the “damage value” of a hit, which in our model is mathematically equivalent to increasing the rate of state transitions).^29,30^

While our approach provides rigorous quantitative insights, we acknowledge several limitations. First, the experimental data is derived from 2D *in vitro* co-cultures. While this simplifies tracking and modeling, it does not fully recapitulate the complex 3D architecture, extracellular matrix barriers, or the immunosuppressive cytokine milieu characteristics of the *in vivo* tumor microenvironment. Consequently, the conjugation rates estimated here may differ in dense tissue where T-cell motility is physically constrained. Second, our ODE framework assumes a well-mixed population (mean-field approximation). Although appropriate for the suspension-like dynamics of the assay, it ignores spatial heterogeneity and local clustering effects that may influence kill rates in a solid tumor context. Finally, while the model identifies that SIN activation enhances the macroscopic killing rate *k*_3_, it remains phenomenological regarding the molecular drivers. The specific downstream signaling pathways (e.g., specific upregulation of Perforin/Granzyme B vs. LFA-1 clustering) triggered by CCL21+ICAM1 interaction require further molecular characterization to fully map the genotype-to-phenotype link.

## Lead Contact

Further information and requests for resources and reagents should be directed to and will be fulfilled by the Lead Contact, Haralampos Hatzikirou or Benjamin Geiger.

## Materials Availability

This study did not generate new unique reagents. The proteins used in this study were produced by the Structural Proteomics Unit at the Weizmann Institute of Science

## Data and Code Availability

- All raw time-lapse microscopy data and the segmented single-cell tracking datasets generated in this study have been deposited and are publicly available as of the date of publication.
- The complete source code for the multi-hit ODE model, including scripts for parameter estimation (profile likelihood) and global sensitivity analysis (Sobol indices), is available on the GitHub page at https://github.com/Mk-guptas/MechanisticModel_Tcell
- Any additional information required to reanalyze the data reported in this paper is available from the lead contact upon request.

## 4 ACKNOWLEDGMENTS

H.H., B.G., and M.K.G. would like to thank Volkswagenstiftung for its support of the “Life?” program (96732). HH has received funding from the Bundes Ministeriums für Bildung und Forschung under grant agreement No. 031L0237C (MiEDGE project/ ERACOSYSMED). HH acknowledges the support of the RIG-2023-051 grant from Khalifa University and the UAE-NIH Collaborative Research grant AJF-NIH-25-KU.

## 5 AUTHOR CONTRIBUTIONS

Conceptualization, H.H. and B.G.; Methodology, M.K.G.,R.Z. and J.S.; Investigation, M.K.G., R.Z., and S.Y.; Formal Analysis, M.K.G. and R.Z.; Visualization, M.K.G. and R.Z.; Writing – Original Draft, M.K.G.; Writing – Review & Editing, all authors; Supervision, B.G. and H.H.; Funding Acquisition, B.G. and H.H.

## STAR ⋆ METHODS

- EXPERIMENTAL MODEL AND SUBJECT DETAILS
- METHOD DETAILS
  - Substrate Functionalization
  - CD8^+^ T Cell Isolation, Activation, and Culture
  - In vitro Cytotoxic T-Cell Killing Assay
  - Cell Segmentation and Image Analysis
  - Multi-hit ODE model
- QUANTIFICATION AND STATISTICAL ANALYSIS
  - Parameter Estimation Strategy
  - Model Selection (AIC)
  - Profile Likelihood Analysis
  - Global Sensitivity Analysis
  - Statistical Tests

## 7 SUPPLEMENTAL INFORMATION

Supplemental information can be found with this link SUPPLEMENTAL INFORMATION

## STAR ⋆ METHODS

### Experimental Model and Subject Details

#### Mice

T-cell receptor (TCR)-transgenic OT-I mice (harboring ovalbumin (OVA)-specific CD8^+^ T cells) were purchased from Jackson Laboratories (Bar Harbor, ME, USA) and bred in the animal facility of the Weizmann Institute. For all experiments, female mice between the ages of 6 and 12 weeks were used. Experiments were performed under protocols approved by the Weizmann Institute’s Institutional Animal Care and Use Committee.

### Method Details

#### Substrate Functionalization

Substrate functionalization was performed by overnight incubation in phosphate-buffered saline (PBS) containing 10 µg/ml CCL21 and 100 µg/ml ICAM1 (produced by the Structural Proteomics Unit of the Weizmann Institute, Rehovot, Israel) at 37°C. Detailed information on the protein production and purification can be found in SI note 4.

#### CD8^+^ T Cell Isolation, Activation, and Culture

Naïve CD8^+^ T cells were purified (*>* 95%) from a cell suspension harvested from the crushed spleens of OT-I mice, using a CD8a^+^ T-Cell Isolation Kit and magnetic-associated cell sorting (MACS), according to the manufacturer’s instructions (Miltenyi Biotec, Bergisch Gladbach, Germany). Freshly isolated CD8^+^ T cells were then activated with anti-CD3 and anti-CD28-coated microbeads at 1:1 bead:cell ratio (Miltenyi Biotec) and with 100 U/ml of IL-2 (BioLegend, San Diego, CA, USA). The cells were suspended in RPMI 1640 medium w/o phenol red, supplemented with 10% FBS, 100 U/ml of penicillin, 100mg/ml of streptomycin, 2mM glutamine, 1mM sodium pyruvate, and 50 mM *β*-mercaptoethanol (Biological Industries, Beit Haemek, Israel) (“complete medium”). The cells were further cultured at 37°C in a 96-well plate (250 *µ*L/well) that was either uncoated or precoated with CCL21-ICAM1 SIN. On day 3 poststimulation, the beads were removed using a magnet, and the contents of all the wells were diluted two fold and subsequently distributed into new wells (coated and uncoated) with fresh medium and IL-2. The cells were then cultured until day 7.

#### In vitro Cytotoxic T-Cell Killing Assay

Target B16 cells expressing OVA and GFP (courtesy of Guy Shakhar, Weizmann Institute of Science) were suspended in complete medium. Cells were seeded in a 96-well polystyrene plate (20,000 cells per well) and incubated for 2 hours, to enable their attachment to the substrate. CD8^+^ T cells, activated with CD3/28 microbeads and cultured on ICAM1+CCL21-coated or uncoated surfaces for 7 days, as described above, were subsequently added, on top of the B16 cells, in a ratio of 3:1 (effector: target). The cells were then monitored by time-lapse imaging using a CellDiscoverer 7 microscope (Carl Zeiss Ltd.). Imaging was performed using the TL-oblique contrast method and EGFP channel, using a Plan-Apochromat 20X/0.95 objective and a 1x Tubelens. Illumination was carried out with a TL-LED lamp and LED-Module 470 nm with an excitation wavelength of 488 nm and emission wavelength of 509 nm. Images were acquired using an Axiocam 702 camera (Carl Zeiss Ltd.) at 5 min intervals, for a total period of 48h.

### Cell Segmentation and Image Analysis

For the microscopy-based morphometric analysis of the cytotoxic process, deep learning cell segmentation software, Cellpose^31^, was used for semi-automatic identification of target cells. Target cells were segmented based on their morphology, imaged by the TL oblique channel, starting with the built-in ‘CPx’ model and refined through a custom model trained via annotation, correction, and validation of the training data. To analyze dynamic morphological changes in target cells during the cytotoxic assay, the trained target cell model was applied to frames captured at multiple time points. Segmentation outputs were subsequently corrected and validated using the GFP channel, which exclusively identifies live target cells. The resulting masks were then processed in Fiji software^32^, where a custom macro was used to convert them into ROIs (Regions of Interest) and to overlay these ROIs onto the corresponding GFP channel frames. Finally, various parameters of target cells, including cell area, circularity, and mean GFP intensity, were calculated.

#### Event detection

- **Target cell death**: Defined as the moment a cell underwent morphological rounding followed by complete loss of cytoplasmic GFP signal. Time of death (the interval between rounding up and complete loss of GFP signal) was recorded for each trajectory.
- **Conjugation**: T cells (visible in the brightfield channel) were segmented separately. A conjugation event was defined when a T cell and tumor cell were co-localized for ≥3 consecutive frames (15 minutes). Contact duration was recorded to estimate *k*_2_.

#### Multi-hit ODE model

We modeled the interaction between T cells and tumor cells using a system of ordinary differential equations (ODEs). The tumor cell population is structured into compartments *C*_*i*_ representing cells that have accumulated *i* lethal hits. The system assumes:

- Tumor growth is logistic and restricted to the undamaged state *C*_0_.
- T cell binding (*k*_1_) and unbinding (*k*_2_) are independent of the damage state of tumor cells.
- Hit delivery (*k*_3_) occurs within the complex and transitions the tumor cell to state *C*_*i*+1_.
- The final state *C*_*n*_ is apoptotic and decays at rate *d*.

The rate equations for the tumor states *C*_*i*_ (*i* = 0 … *n*) and the complexes are:

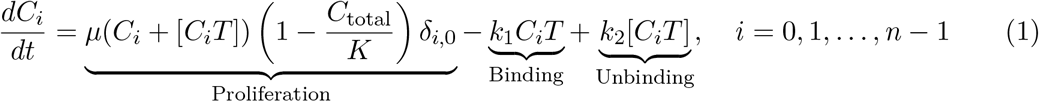

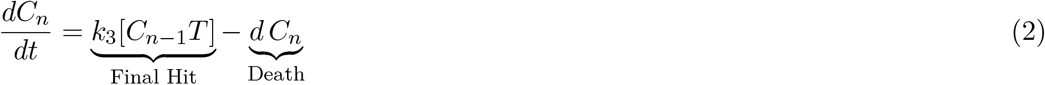

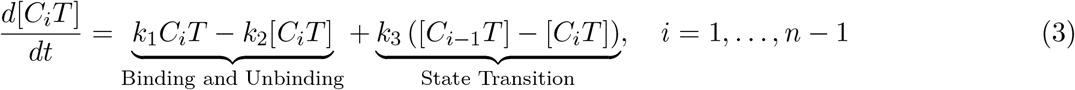

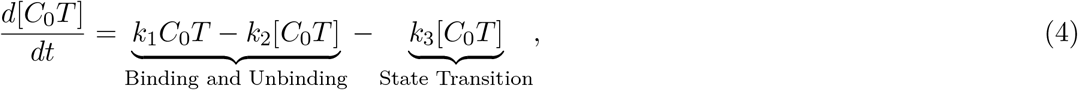

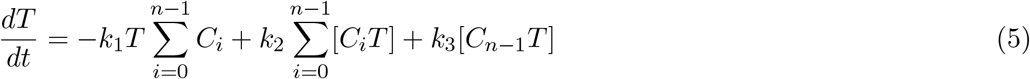

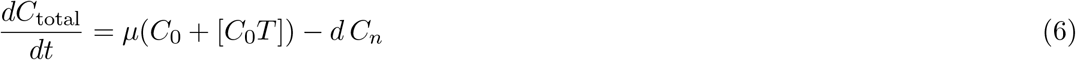

For the final state *n*, the term − *k*_3_ represents the dissociation of the T cell from the dying cell (assuming the T cell leaves to find a new target).

### Quantification and Statistical Analysis

#### Parameter Estimation Strategy

To avoid the identifiability problem, we employed a hierarchical estimation approach.

1. **Direct Estimation (*d, k*_2_):** The decay rate *d* of target cell death was calculated as the inverse of the mean cell death duration (*T*_*death*_) measured from single-cell tracks. The dissociation rate *k*_2_ was calculated as the inverse of the mean effector–target attachment duration (*T*_*attach*_), determined by tracking the time interval for which an individual T cell remained attached to a tumor cell.
2. **Indirect Estimation (*k*_1_, *k*_3_, *µ*):** The remaining parameters were estimated by fitting the ODE model to the population-level time-course data (Total Tumor Count vs. Time). We used the Trust Region Reflective least-squares algorithm (`scipy.optimize.least_squares`) to minimize the objective function:

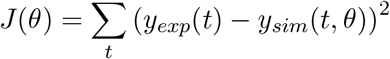

where *θ* = {*k*_1_, *k*_3_, *µ*} is the parameter vector. The carrying capacity in the logistic growth model was set to 500, estimated by dividing the total area of the region of interest by the average projected area of an individual tumor cell.

#### Model Selection (AIC)

To determine the optimal number of hits (*n*), we fitted the model for values of *n* ranging from 1 to 9. We compared the models using the Akaike Information Criterion (AIC):

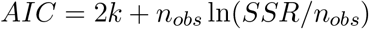

where *k* is the number of free parameters, *n*_*obs*_ is the number of data points, and *SSR* is the sum of squared residuals. The value of *n* that minimized the AIC was selected as the consensus model structure (*n* = 4).

#### Profile Likelihood Analysis

To assess the practical identifiability of *k*_1_ and *k*_3_, we computed the profile likelihood. For a parameter *θ*_*i*_, the profile likelihood 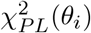 is defined as the minimum *χ*^2^ value (or weighted SSR) obtainable for the model when *θ*_*i*_ is fixed to a specific value and all other parameters *θ*_*j*≠*i*_ are re-optimized. 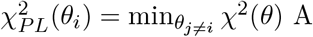 A parameter is considered identifiable at the 95% confidence level if its profile likelihood crosses the threshold Δ_*𝒳*_^2^ = 3.84 (the 95th percentile of the 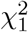 distribution) within the biologically relevant range. Flat profiles indicate non-identifiability. We resolved initial non-identifiability by co-fitting multiple datasets with different initial conditions (E:T ratios). For improved visualization, the profile likelihood is presented in terms of mean squared error (MSE). The derivation of the corresponding ΔMSE threshold for the 95% confidence interval is provided in the SI Note 1.

#### Global Sensitivity Analysis

We performed a variance-based global sensitivity analysis using first-order Sobol indices. As the model output, we considered the area under the curve (AUC) of the tumor population trajectory over 48 hours. The parameter space was sampled using quasi-random Sobol sequences (*N* = 10,000 samples), with parameters varied independently over the following ranges: *k*_1_ ∈ [5.5 × 10^−6^, 5.5 × 10^−5^], *k*_2_ ∈ [10^−3^, 10^−2^], *k*_3_ ∈ [10^−3^, 10^−2^], *µ* ∈ [10^−4^, 10^− 3^], and *d* ∈ [10^− 3^, 10^− 2^]. We then computed the first-order Sobol index for each parameter, which quantifies its main effect on the variance of the model output while excluding higher-order interaction effects.

#### Statistical Tests

Differences in estimated parameter distributions between SIN-treated and untreated groups were assessed using an approximate Wald-type z-test based on profile-likelihood-derived confidence intervals (SI Note 3).

## Supplementary information

### SI Note 1 : Confidence r egions f rom profile statistics

#### Using the 𝒳^2^ profile

For a m odel w ith *i ndependent G aussian e rrors* o f (known o r pooled) variance *σ*^2^, let

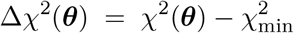

be the likelihood-ratio statistic obtained by fixing the *q* parameters of i nterest at ***θ*** and reoptimising all nuisance parameters. Wilks’ theorem implies that under the null hypothesis “***θ*** equals the true value” Δ_*𝒳*_^2^ is asymptotically 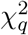 -distributed. Hence a simultaneous two-sided 100(1 ^−^ *α*) % confidence region for the *q* profiled parameters is

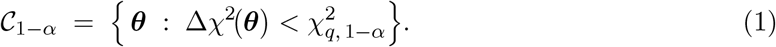

#### Converting the rule to an MSE profile

If the vertical axis of the profile plot is the *mean-squared error*

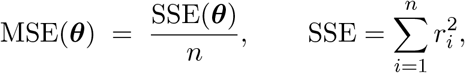

the link between the two statistics is simply

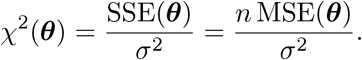

Therefore the horizontal threshold that encloses the same 100(1 − *α*) % confidence region becomes

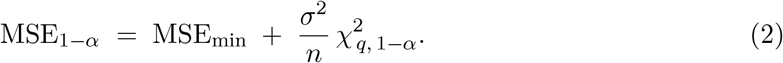

#### Computation of threshold

To establish the horizontal threshold corresponding to the 95% confidence interval in the MSE profile plots, an estimate of the residual variance *σ*^2^ is required (Eq. 2). Rather than treating the variance as a point-wise quantity, we estimated *σ*^2^ as the mean of the individual per-observation variances computed across all available experimental data points. This pooling approach yields a single, common variance estimate that is more statistically robust than any individual point estimate and is consistent with the assumption of homoscedastic errors underlying the least-squares framework. The resulting pooled estimate was *σ*^2^ = 100 (in units of cells^2^). Furthermore, since parameter estimation was performed by co-fitting two datasets simultaneously—each comprising 16 time points—the total number of observations entering the objective function was *n* = 32. Substituting these values into Eq. 2, with *q* = 1 degree of freedom (corresponding to the profiling of a single parameter at a time) and the 95th percentile of the 𝒳^2^ distribution 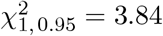, the threshold increment is:

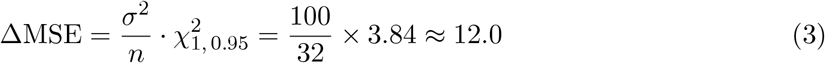

Accordingly, the 95% confidence interval for each parameter is defined as the contiguous set of parameter values for which the profile MSE does not exceed MSE_min_ + 12.0, where MSE_min_ is the minimum value attained along the profile curve.

### SI Note 2: Identifiability test

**SI Fig. 1.**
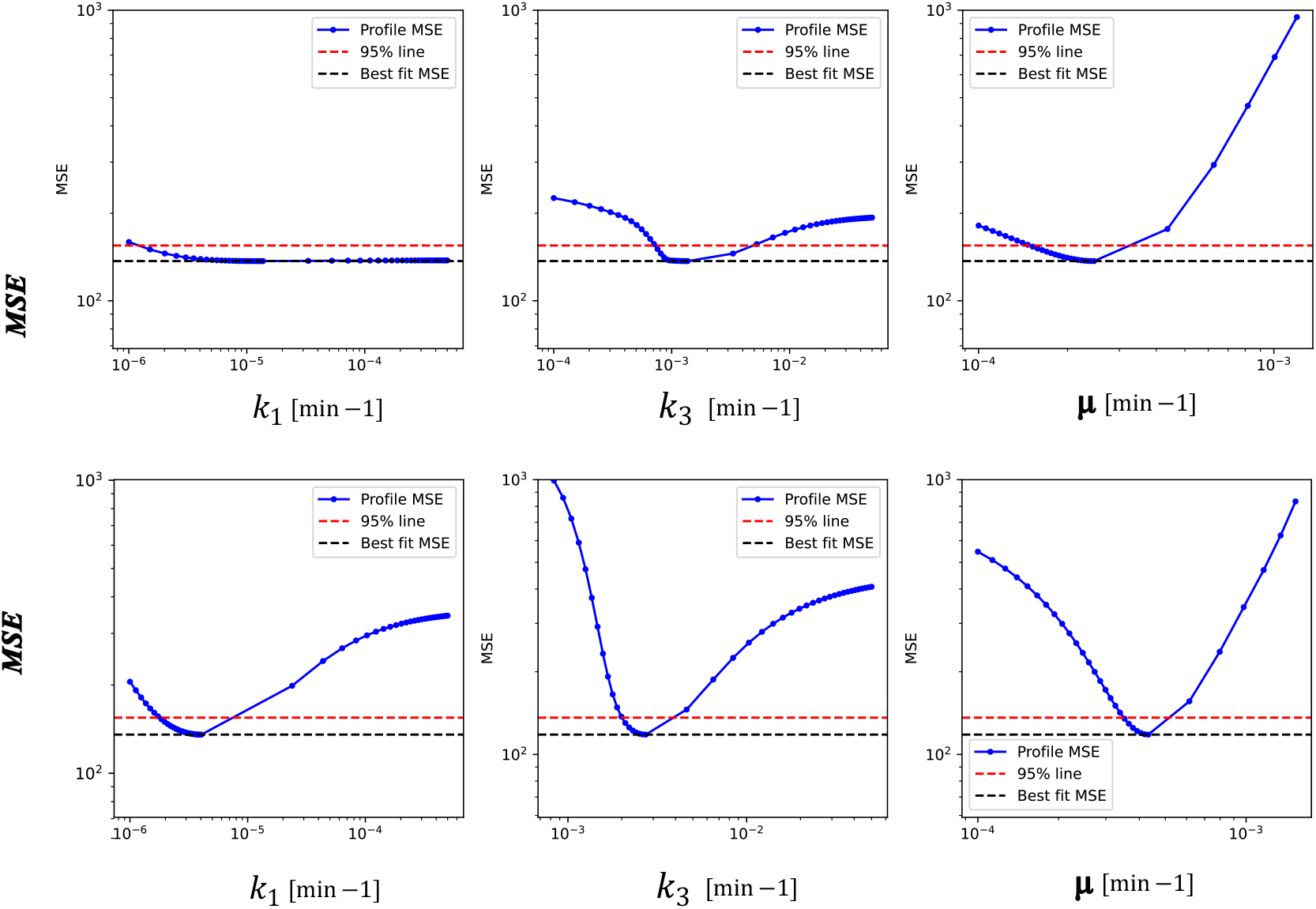
Simulation-based practical identifiability analysis using profile likelihood plots. The upper panels show the analysis performed with a single dataset, where parameters *k*_1_ and *k*_3_ exhibit wide confidence intervals and high standard deviations, indicating poor identifiability. In contrast, the lower panels present the analysis after incorporating an additional dataset, which narrows the confidence intervals and renders *k*_1_ and *k*_3_ identifiable. The solid blue curves represent the sum of squared residuals (SSQ) as each parameter is varied, while the dashed lines denote the best-fit SSQ values.

### SI Note 3: Approximate comparison of parameter estimates between SIN-treated and untreated conditions

To assess whether the fitted parameter values differed between SIN-treated and untreated conditions, we performed an *approximate two-sided Wald z-test* for the null hypothesis

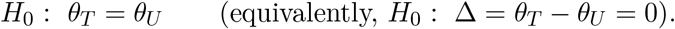

Let 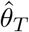 and 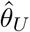 denote the parameter estimates obtained from the treated and untreated datasets, with corresponding standard errors *SE*_*T*_ and *SE*_*U*_ . Under a large-sample (local normality) approximation, we assume

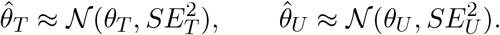

Assuming the two estimates are independent, the difference 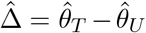 is approximately normal with

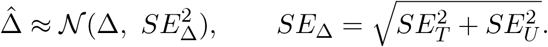

We therefore form the Wald statistic

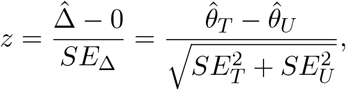

which is approximately standard normal under *H*_0_ (i.e., *z* ≈ 𝒩 (0, 1)). The corresponding two-sided *p*-value is computed as

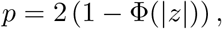

where Φ(·) is the cumulative distribution function of the standard normal distribution.

#### Approximation of standard errors from reported 95% confidence intervals

Because only the estimated means and their 95% confidence intervals were available, the standard errors were approximated from the reported interval widths using the normal approximation

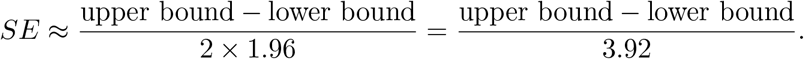

This approximation is exact only for symmetric confidence intervals derived from a normal approximation, and should therefore be interpreted cautiously when the reported intervals are asymmetric.

#### Statistical test for *k*_3_

For *k*_3_, the reported values were

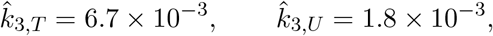

with 95% confidence intervals

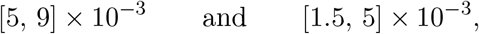

for treated and untreated conditions, respectively. Therefore,

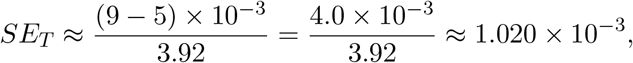

and

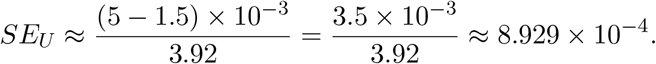

Thus, the standard error of the difference is

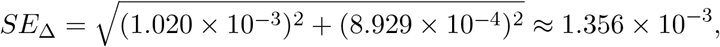

and the difference in means is

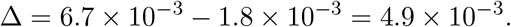

Hence,

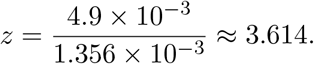

The corresponding two-sided *p*-value is

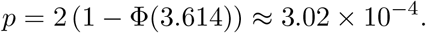

Under this approximation, the difference in *k*_3_ between treated and untreated conditions is statistically significant at the 5% level.

#### Statistical test for *µ*

For *µ*, the reported values were

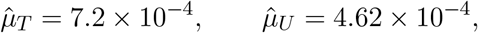

with 95% confidence intervals

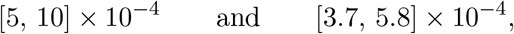

for treated and untreated conditions, respectively. Therefore,

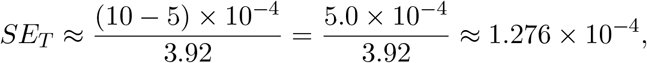

and

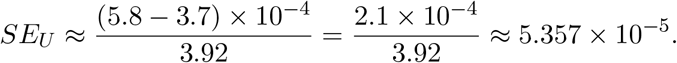

Thus,

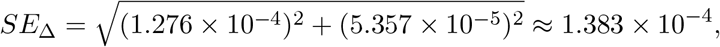

while the difference in means is

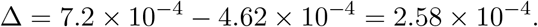

Hence,

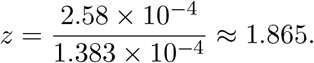

The corresponding two-sided *p*-value is

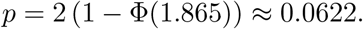

Under this approximation, the difference in *µ* between treated and untreated conditions is not statistically significant at the 5% level.

These *p*-values should be interpreted as approximate, since they rely on a local Gaussian approximation and on standard errors inferred from reported confidence interval widths. In particular, when confidence intervals are asymmetric or derived from profile likelihood, a likelihood-ratio or profile-based comparison is generally more reliable than the Wald approximation.

### SI Note 3: Condition-dependent contact time and death duration

**SI Fig 2:**
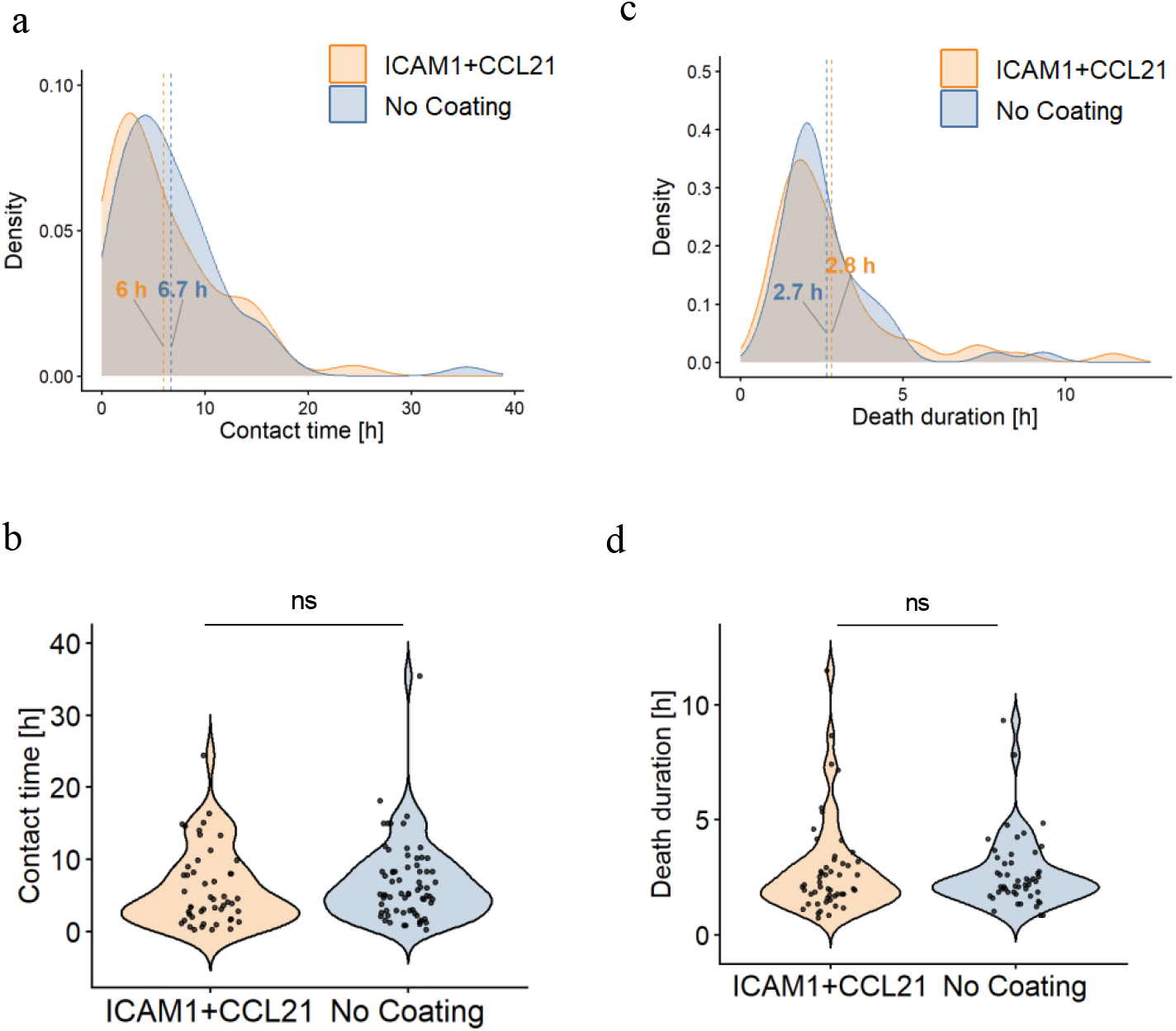
Experimental estimate of dissociation rate (k_2_) and cell death decay rates (d) for SIN-treated and untreated T cells. **(a and b)** Density plots of the contact time between target and T cells by pretreatment. The vertical line in **(a)** indicates the mean. Statistical significance was assessed using Welch’s t-test (α = 0.05; ns, not significant). **(c and d)** Density plots of the interval between target cell rounding and GFP loss (“Cell Death duration”) by pretreatment. The vertical line in **(c)** indicates the mean. Statistical significance was assessed using Welch’s t-test (α = 0.05; ns, not significant).

### SI Note 4: Protein production and purification

#### Production and purification of mouse CCl21 (mCCL21)

The gene encoding mCCL21 (24–133) followed by a TEV cleavage site, a 6 x Proline rigid linker and a C-terminal 8 x His tag was optimized for *E. coli* expression and cloned into pET21a. The resulting construct was transformed into *E. coli* BL21 (DE3) competent cells. Bacterial precultures were grown overnight at 37 ^°^C with shaking and used to inoculate 7.5 L LB. Cultures were grown at 37 ^°^C with shaking to an optical density OD_600_ = 0.6. Protein expression was induced by the addition of isopropyl-*β*-D-thiogalactopyranoside (IPTG; 200 µM), and the culture was kept at 15 ^°^C with shaking overnight. Cells were harvested by centrifugation, and the pellet was resuspended in 100 ml lysis buffer containing PBS, 150 mM KCl, 10 mM imidazole, 1 mM DTT supplemented with 0.2 mg/ml lysozyme, 20 µg/ml DNase, 1 mM MgCl_2_, and protease inhibitor cocktail (Calbiochem set 3). After lysis by a cooled protein disrupter (Constant Systems), the soluble fraction was obtained by centrifugation and purified by immobilized metal ion affinity chromatography (IMAC) using a HiTrap FF_5 ml cartridge (Cytiva) using an FPLC system (ÄKTA GE Healthcare Life Sciences) at 4 ^°^C. The column was washed with lysis buffer containing 50 mM imidazole and mCCL21 was eluted in one step with elution buffer (PBS, 150 mM KCl, 500 mM imidazole, 1 mM DTT). The fractions containing mCCL21 were analyzed using SDS-PAGE, pooled and applied to an Xbridge C4 (Waters) reverse phase column operated by an HPLC system (Waters) pre-equilibrated with binding buffer containing 5% acetonitrille, 0.1% TFA in DDW. The protein was eluted with a linear gradient to 95% acetonitrille, 0.1% TFA in DDW. Pure mCCL21 samples from consecutive runs were pooled and lyophilize to dryness. The dry protein was resuspended in PBS and divided into small aliquots before flash freezing in liquid nitrogen. mCCL21 was typically characterized by SDS-PAGE and WB (developed with anti-His).

##### Expression and purification of mouse ICAM1 (mICAM1)

mICAM1 (aa residues 1–485) followed by a TEV recognition site, a 6 x Proline rigid linker and an 8 x His tag was cloned into the pHLSec vector for secretion from Expi293 human cells. Medium (400 ml) containing the secreted protein was collected 72 h post-transfection for purification. After carefully removing the cells, the medium was applied to a Ni column (HisTrap FF_ 5 ml) equilibrated with PBS and the bound mICAM1 was washed with PBS containing 50 mM imidazole and eluted with PBS containing 500 mM imidazole. Fractions containing mICAM1 were pooled, concentrated and injected onto a Superdex_200_16/60 PrepGrad size exclusion column equilibrated with PBS. The pure mICAM1 migrated at 77 ml as a single peak. The identity of the protein was confirmed by SDS-PAGE developed with Coomassie blue and WB developed with anti-His. Aliquots were flash-frozen with liquid nitrogen and kept at ^−^80 ^°^C.

